# The evolution of dengue-2 viruses in Malindi, Kenya and greater East Africa: epidemiological and immunological implications

**DOI:** 10.1101/850107

**Authors:** S Pollett, K Gathii, K Figueroa, W Rutvisuttinunt, A Srikanth, JN Nyataya, BK Mutai, G Awinda, RG Jarman, I Maljkovic Berry, JN Waitumbi

## Abstract

Kenya experiences a substantial burden of dengue, yet there are very few DENV-2 sequence data available from this country and indeed the entire continent of Africa. We therefore undertook whole genome sequencing and evolutionary analysis of fourteen dengue virus (DENV)-2 strains sampled from Malindi sub-County Hospital during the 2017 DENV-2 outbreak in the Kenyan coast. We further performed an extended East African phylogenetic analysis, which leveraged 26 complete African *env* genes. Maximum likelihood analysis showed that the 2017 outbreak was due to the Cosmopolitan genotype, indicating that this has been the only confirmed human DENV-2 genotype circulating in Africa to date. Phylogeographic analyses indicated transmission of DENV-2 viruses between East Africa and South/South-West Asia. Time-scaled genealogies show that DENV-2 viruses are spatially structured within Kenya, with a time-to-most-common-recent ancestor analysis indicating that these DENV-2 strains were circulating for up to 5.38 years in Kenya before detection in the 2017 Malindi outbreak. Selection pressure analyses indicated sampled Kenyan DENV strains uniquely being under positive selection at 6 sites, predominantly across the non-structural genes, and epitope prediction analyses showed that one of these sites corresponds to a putative predicted MHC-I CD8+ DENV-2 Cosmopolitan virus epitope only evident in a sampled Kenyan virus. Taken together, our findings indicate that the 2017 Malindi DENV-2 outbreak arose from a strain which had circulated for several years in Kenya before recent detection, has experienced diversifying selection pressure, and may contain new putative immunogens relevant to vaccine design. These findings prompt further genomic epidemiology studies in this and other Kenyan locations to further elucidate the transmission dynamics of DENV in this region.

**Author summary (non-technical):** Kenya experiences a substantial burden of dengue, yet the patterns of dengue spread in this region are unclear. Evolutionary analyses of dengue virus strain sequences could offer major insights into the spread of dengue viruses in this region, but there are very few DENV-2 sequence data available from this country and indeed the entire continent of Africa. We therefore undertook whole genome sequencing and evolutionary analysis of fourteen dengue virus (DENV)-2 strains sampled from Malindi sub-County Hospital during the 2017 DENV-2 outbreak in the Kenyan coast. We further performed an extended East African phylogenetic analysis, which leveraged 26 complete African *env* genes. Our results indicated transmission of DENV-2 viruses between East Africa and South/South-West Asia, that there is localized spread of DENV-2 viruses within Kenya, and that Kenyan DENV-2 strains were circulating for up to 5.38 years in Kenya before detection in the 2017 Coastal outbreak. We further performed selection pressure analyses and identified possible new markers of DENV-2 immune recognition specific to this population, with relevance to vaccine design.

## Introduction

Dengue poses a persistent and ongoing threat to public health in many tropical countries, with an estimated global burden exceeding half a billion infections per year [1]. In hyper-endemic countries, the annual force-of-infection is 5-10%, resulting in the majority of the population exposed by adulthood [2, 3]. Non- or pauci-immune populations may also experience intermitted outbreaks with attack rates exceeding 50% [4]. In addition to an increased number of dengue endemic countries in the last three decades, some dengue hyperendemic countries have seen up to a 500% increase in dengue mortality over a 15 year period [5]. Dengue is caused by a serogroup of four dengue viruses (DENV). These are single positive-strand RNA viruses that belong to the *Flaviviridae* family, and are borne most effectively by the *Aedes aegypti* mosquito [6]. The escalating burden of dengue is in part due to the rapid urbanization of tropical regions, and in part due to the incomplete cross protection offered by a primary dengue infection [7]. Current countermeasures for dengue include vector control and optimized medical management following established guidelines. There is currently only one licensed dengue vaccine (Dengvaxia), which has been implemented in Brazil and the Philippines, and has major safety and effectiveness concerns in dengue unexposed populations [8].

East Africa is thought to have played a key role in the global spread of DENV in the 18th century, with apparent dengue case reports described there as early as 1823. There is a popular – but unproven - hypothesis that dengue was introduced into the Caribbean from East Africa [9, 10]. Indeed, the very term ‘dengue’, is believed to have been derived from the Swahili language (then called ‘dinga’) [10]. However, compared to other tropical areas, the epidemiology of dengue in East Africa, and Africa in general, is not well understood [7]. This is attributed to limited surveillance and limited availability of diagnostics for dengue, a disease that often has similar clinical symptoms and signs as other endemic tropical diseases in Africa, such as malaria [11, 12]. Further, there are immense competing public health demands in many countries from this region [12]. Nevertheless, the burden of dengue in East Africa is being increasingly recognized. Studies have indicated the circulation of DENV in Kenya as well as Djibouti, Eritrea, Ethiopia, Mozambique, Somalia, Sudan, Tanzania, and Uganda [9, 13–18].

Kenya, in particular, has experienced substantial DENV outbreaks in coastal regions since at least 1982 when DENV-2 was first detected in the towns of Malindi and Kilifi [19, 20]. Since the initial detection in the early 1980s, a substantial DENV-2 outbreak occurred in the North-Eastern town of Mandera in 2011, which was thought to be introduced from Somalia based on its proximity to the Kenyan border [21]. A substantial DENV outbreak was noted in the urban Mombasa in 2013, with an attack rate of approximately 13% in one district alone [22]. Extended dengue testing on febrile cases presenting to a range of hospitals throughout Kenya in 2011-2014 also indicated the circulation of DENV serotypes 1 - 3 in the coastal locations of Mombasa, Malindi and Lamu, the North-Eastern locales of Garissa and Wajir, and the Western capital of Nairobi [23]. More recently, through molecular characterization of undifferentiated febrile cases, DENV 1 −4 was detected in a 2014 - 2015 pediatric febrile surveillance cohort in Western Kenya (Chulaimbo and Kisumu), although it remains unclear why dengue outbreaks are more frequently noted on the Kenyan coast [24].

The population seroprevalence of DENV in Kenya is highest in coastal regions, where DENV seropositivity has exceeded 58.8% across all ages tested, and has exceeded 20% in the under ten year olds [25]. Western Kenyan seroprevalence is estimated to be considerably lower, although still substantial in pediatric age strata, suggesting inter-epidemic transmissions occurs in both Western and Coastal Kenya [26]. *Aedes aegypti* is prevalent in both coastal Kenya as well as more Western regions, including the Kisumu and the capital Nairobi [12, 27–29]. Ecological and demographic changes, such as increasing irrigation farming, international travel, and urbanization, are believed to be risk factors for ongoing dengue outbreaks in this country [12, 22, 30].

Despite the increasingly recognized burden of DENV in Kenya, there has been few DENV sequence data published from this country. The 2011-2014 Mombasa outbreak yielded short partial *env* sequences, which only permitted genotyping [23]. The paucity of sequence data from Kenya has limited our understanding of the epidemic dynamics of recent Kenyan DENV outbreaks. Further, there is a lack of DENV African genomic data in general. A recent comprehensive assessment of the global adequacy of dengue genetic sampling indicated that Africa represents just 1% of all published DENV full *env* sequence data [31]. In the case of DENV-2, there are very few African-sampled whole genome DENV-2 sequences published. This lack of African DENV sequence data has long confounded attempts to elucidate the role of East Africa in global DENV spread, and has limited any inference about whether recently developed and/or licensed dengue vaccine products may be mismatched to circulating dengue viruses in Africa [32–34].

To redress this dearth of Kenyan DENV genomic data, and to investigate a recent Kenyan DENV-2 outbreak within a phylodynamic framework, we undertook whole genome sequencing and evolutionary analysis of 14 DENV-2 whole genomes. These data represent a major increase to the previously available number of African DENV-2 whole genome data. Specifically, we sought to estimate the duration of cryptic circulation of DENV-2 before first case detections in the 2017 in Malindi outbreak and determine whether this outbreak could be partly explained by viral immune escape. Leveraging these new data and all existing public DENV data, we also performed an extended regional East African molecular epidemiology of 26 complete African *env* genes to offer insight into dispersal and population structure of DENV-2 viruses in East Africa relative to other African and other global regions. In order to support future vaccine development and evaluation, we further explored whether there were any novel T cell epitopes identified along the DENV-2 whole proteome relevant to the Kenyan population.

## Methods

### Field methods: study population, setting and dengue case detection

The coastal city of Malindi, Kenya, is located 120 kilometers Northeast of Mombasa, a major urban seaport of Kenya. Malindi is the largest urban area of Kilifi County, with a population of 207,000 persons [35], and is known to be endemic for dengue [23]. Patients with suspected dengue cases presenting to Malindi sub-county Hospital, Malindi, Kilifi County in 2017 for clinical care were consented under an IRB approved protocol that recruits patients with acute febrile illness. Dengue infections were confirmed in 14 suspect cases by genotyping using EasyScreen flavivirus PCR typing kit (Genetic Signatures, Australia).

### RNA extraction and sequencing methods at the Basic Science Lab (Walter Reed Project/KEMRI, Kisumu, Kenya)

From the 14 confirmed DENV-2 infections, RNA extraction and sequencing was performed at the Basic Science Lab (KEMRI, Kisumu, Kenya as recently described by Gathii et al [19]. Briefly, total RNA was extracted from the sera using the Direct-zol miniprep kit (Zymo Research). cDNA synthesis was performed by sequence-independent single-primer amplification following methods by Djikeng et al [36], followed by cDNA amplification using MyTaq DNA polymerase (Bioline, MA). The Nextera XT kit (Illumina, San Diego, CA) was used to prepare sequence libraries before sequencing on the MiSeq platform (Illumina, San Diego, CA). Their sequence data underwent assembly and annotation using CLC Genomics Workbench version 8.5.1 (Qiagen), using a DENV-2 reference genome (GenBank accession number: NC001474). Whole genome sequences were derived from 10 of the 14 specimens (GenBank accession numbers: MG779194 - MG779203.

### RNA extraction and sequencing methods at the Department of Virology (WRAIR)

Extracted RNA from an additional four DENV-2 infected sera that failed to sequence in Kenya underwent DENV-2 whole genome sequencing at the Walter Reed Army Institute of Research using in-house designed primers which were optimized to the Kenyan DENV-2 sequence data generated by Gathii et al. Amplicons were generated from extracted RNA by these DENV-2 specific primers in addition to random primers. A conventional amplification using DENV-2 specific primers (Table S1) were generated with Taq polymerase (ThermoFisher, Waltham, MA). Random amplicons were obtained using SuperScript III RT and HiFi Taq (ThermoFisher, Waltham, MA). Enhanced sensitivity in amplification was pursued utilizing microfluidic amplification with DENV-2 specific and random primers and SSIII/HiFi Platinum Taq on the integrated fluidic circuits (IFC) (Fluidigm, Palo Alto, CA). The reaction conditions for both conventional and microfluidic amplifications were 50°C for 30 minutes and 94°C for 2 minutes followed by 35 cycles of 94°C (30 seconds), 55°C (30 seconds), and 68°C (2 minutes), and a hold at 68°C for 7 minutes prior to cooling down to 4°C. Amplicons were used for NexteraXT libraries (Illumina, San Diego, CA) and QIASeq Fx libraries (QIAGEN, Germantown, MD). Libraries were validated using Qubit (ThermoFisher, Waltham, MA) and TapeStation (Agilent, Santa Clara, CA). The sequencing was conducted on the MiSeq reagent v.3 600 cycles (Illumina, San Diego, CA). The four genomes were assembled using ngs_mapper v1.5, an in-house reference mapping pipeline [37], and curated manually for removal of sequencing and assembly-associated errors. Ngs_mapper output files, VCF, BAM and statistical read quality/depth graphs, were used to support base-call curations. Pipeline parameters, minimum base quality and allele frequency thresholds, were kept at respective defaults of Q25 and 20% for consensus sequence reconstruction. The accession number of these genomes sequenced at WRAIR were MN335244-MN335247.

#### Evolutionary analysis

##### Genotyping, recombination detection and phylogenomic analysis 14 Kenyan DENV strains

All 14 full genome sequences (including the 10 full genomes recently reported by Gathii et al) were collated with a comprehensive curated reference dataset of 1212 published annotated whole genome DENV-2 data available on the VIPR and NCBI Genbank databases as of April 2018 [38, 39] (Table S2).

Alignment was performed using MAFFT and manually edited thereafter, before truncation to coding regions [40]. Initial neighbor-joining trees were used to confirm the genotype of these data as the Cosmopolitan lineage using MEGA version 7.0 [41]. Thereafter, we inferred a maximum likelihood tree using all new Kenyan whole genome data, all published Cosmopolitan lineage whole genome data (n = 274), and an American genotype outgroup strain (GenBank accession number HM582104). This dataset included the three African DENV-2 Cosmopolitan genomes available at the time of GenBank search, including n = 1 from Tanzania, n = 2 from Burkina Faso. Notably, this whole genome analysis excludes the 200 nt partial env-gene sequences derived from the 2011-2014 Mombasa outbreak [23], as such short sequence lengths lack appropriate phylogenetic resolution. We screened for recombinants using the RDP4 program [42], as well as screening for incongruences in neighbor-joining trees of four subgenomic regions (CDS nucleotide positions 1-2543, 2544-5087, 5088-7630 and 7631-10173) inferred using MEGA 7.0. We removed those sequences with suspected recombination. A final maximum likelihood phylogeny was inferred using the PhyML software using the NNI and SPI tree search method and the aLRT approach for node support [43]. A GTR+I+G nucleotide substitution model was selected by Akaike information criterion (AIC) criteria using JModelTest2 [44].

##### East African molecular epidemiology analysis leveraging an extended env-gene dataset

Due to major whole genome sampling gaps in Kenya, and indeed throughout Africa, we also inferred an *env*-gene tree, including more sequences, to estimate the regional and international phylogeography of DENV-2 in East Africa. This leveraged an extended Cosmopolitan genotype complete *env*-gene alignment (Table S3) containing n = 1343 complete *env* sequences, including all African full-length env sequence data available at the time of GenBank search (n = 26 total, including Burkina Faso n = 4, Somalia n = 6, Tanzania n = 1, Uganda n = 1 and the Kenyan data from this study n = 14). We inferred a maximum likelihood tree using the PhyML software, with a GTR+I+G nucleotide substitution model selected by AIC in jModelTest2 [44].

##### Time-scaled Bayesian phylogeographic analyses to reconstruct the history of the Kenyan 2017 DENV outbreak

All 14 genomes were analyzed using Bayesian phylogeography to further clarify the time-scale and rates of Kenyan DENV virus evolution. A whole-genome time-scaled phylogeny was inferred using the BEAST package [45]. Given the large computational load of performing Bayesian analyses on a large serotype-wide full-genome dataset, we restricted the analysis to strains comprising the basal sublineage of the Cosmopolitan genotype which contained all Malindi Kenyan data (Fig 1). Root-to-tip regression was performed using Temp-est which indicated adequate temporal structure of this subsampled data (coefficient of determination = 0.91, n = 37 sequences). Time-scaled Bayesian analysis was performed using a nucleotide model of SYM+I+G as selected using AIC criteria in jModelTest2. A maximum clade credibility tree was inferred using the posterior distribution of 10,000 trees sampled from a 200 million Markov Chain under a relaxed clock assumption (uncorrelated lognormal distribution) and a skyline demographic model. Statistical convergence was determined by examining trace plots in addition to ESS values, which exceeded 200 for all parameters.

**Figure 1.**
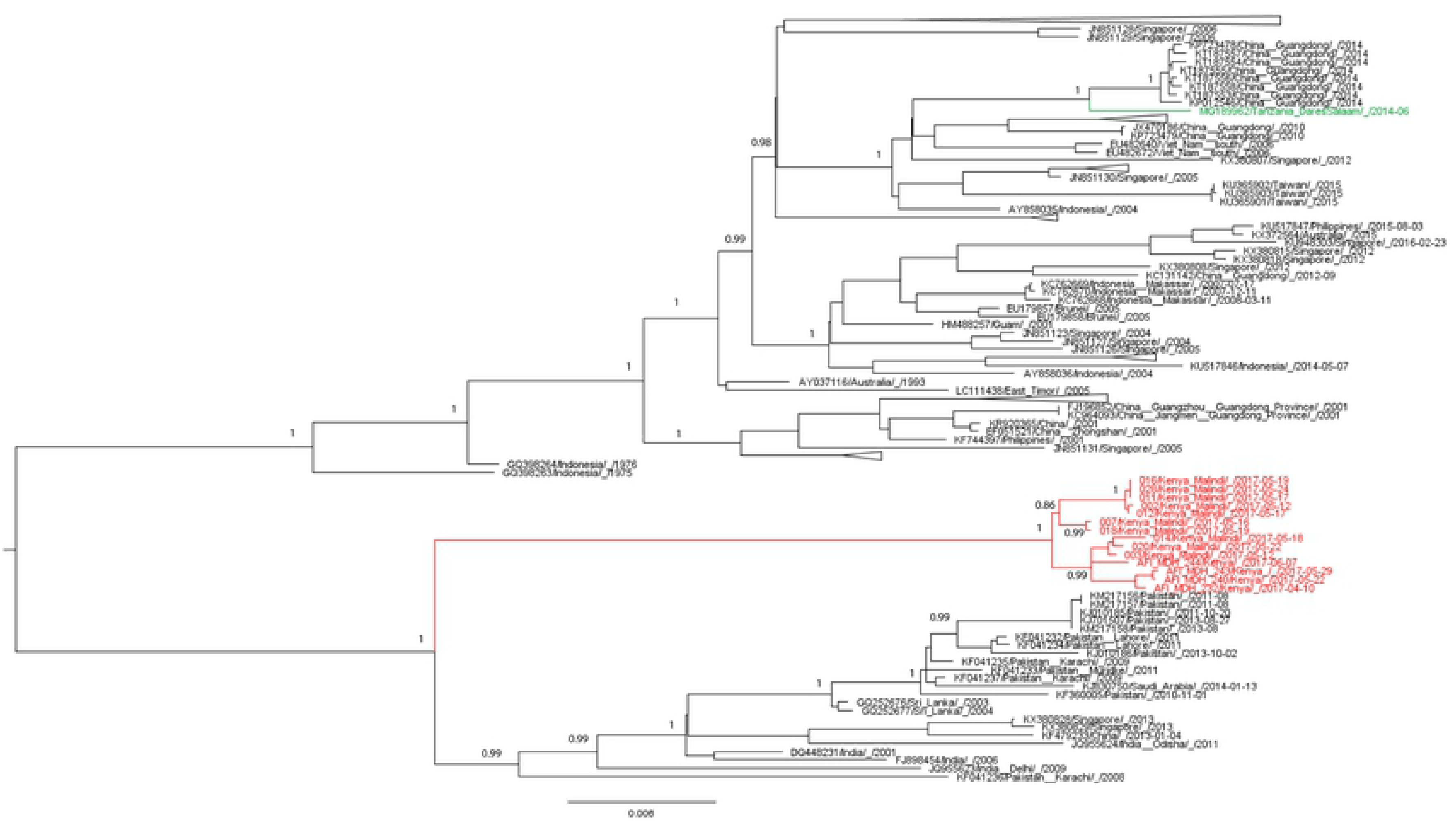
Whole genome maximum likelihood phylogenetic tree of DENV2 Cosmopolitan genotype. American genotype outgroup strain (HM582104/American_Somoa/1972) has been removed for clarity. Numbers indicated aLRT support for key nodes, with aLRT values ≥ 0.75 indicating robust support. Kenyan genomes are noted in red, Tanzania data is indicated in green, all other countries are represented by black. Some well-supported clades were collapsed for clarity. Scale bar indicates genetic distance substitutions/site.

##### Selection pressure analysis

We estimated gene-wide and site-specific selection pressure across the entire genome of all Malindi Kenyan and publicly available unique DENV-2 Cosmopolitan strains in addition to a single American genotype outgroup strain (n = 275 taxa). The SLAC, FUBAR and FEL methods available in the HyPhy package were employed [46, 47]. We used multiple selection pressure detection methods as they substantially vary in sensitivity and specificity [7]. The computational demand of selection pressure analysis across a whole genome dataset of this size required separate analyses on three partitions of the DENV-2 genome: the structural genes (C-prM-env), non-structural genes NS1 through NS3, and non-structural genes NS4 & NS5.

##### MHC-I CD8+ epitope prediction within DENV-2 Cosmopolitan proteomes from Kenya and beyond

CTL (Cytotoxic T cell Lymphocyte) epitope discovery, particularly in critically under-sampled regions like Africa, has major relevance for dengue vaccine development and evaluation. These Kenyan data, compiled with a comprehensive global DENV-2 Cosmopolitan genotype curated dataset, allowed us to examine whether strains from this global region may contain possible novel CTL epitopes. We first predicted the HLA MHC-I haplotypes of the hosts sampled in this study using a population genetics framework, which leveraged the curated www.allelefrequencies.net database. The C*06:02, C*04:01 and A*02:01 alleles were selected based on their relatively high prevalence across all published Kenyan HLA admixture studies and because they are considered to be relatively frequent alleles in the global human population [48]. Further, the HLA A*02 haplotype is an HLA super-type with high MHC-I cross reactivity [49].

Using these selected HLA molecules we performed MHC-I affinity prediction across sequential 9mer peptides across the entire DENV-2 proteome of all Kenyan whole genome data using the Artificial Neural Network method [50]. We then determined whether any of these predicted 9mer peptides correspond to previously *in-vitro* studied linear CD8+ or CD4+ epitopes in the IEDB database [51]. Finally, we performed a conservancy analysis of these 9mer peptides to determine whether such putative epitopes are possibly geographically restricted.

##### Ethics

Field methods, laboratory and viral genomic analyses of dengue positive cases was performed in concordance with the Ethical Review Committee of the Kenya Medical Research Institute (SSC protocol #1282) as well as the Human Subject Protection Branch of the Walter Reed Army Research Institute (WRAIR protocol #1402). All participants gave informed consent. A parent or guardian of any child participant provided informed consent on the child’s behalf. In addition, children between 13 and <18 years provided assent to participate. Consent was written if participant was literate and fingerprint if illiterate, with the signature of an independent witness.

## Results

### Clinical and demographic characteristics

All the 14 patients from whom the whole DENV genome sequences were derived were recruited in the months of April to June 2017. All had typical signs of dengue fever comprising high fever (>38.5 °C), headache, chills, muscle and joint pains and body rash.

#### Kenyan DENV-2 belongs to the Cosmopolitan genotype and there is no evidence of DENV-2 recombinants circulating in Kenya

Genotype determination showed that these DENV-2 strains sampled from Kenya, and indeed all other East African countries sampled to date, are of the Cosmopolitan genotype. None of the Kenyan genomes were found to be recombinant, although there was significant (p<0.05) recombination signal detected in a Sri Lanka genome (accession number: FJ882602) by 2 of 7 methods used in RDP4 (RDP and BootScan), indicating a ~700bp insert from a parental genome detected to be Pakistan_Karachi/2008 (accession number: KF041236). As this constitutes borderline evidence of recombination, we further tested for phylogenetic incongruence by dividing the full genome alignment into 4 equal length alignments. NJ phylogenetic trees (Fig S1) were constructed for each part of the genome to assess changes in tree topology along the genome, which was indeed noted. Because phylogenetic inference is affected by recombination, the Sri Lanka 2008 genome was removed from the alignment [7]. This approach also identified that the position of the two Burkina Faso human cosmopolitan strains (accession numbers: EU056810, GU131843), and a further five Indonesian whole genomes (accession numbers: GQ398260, GQ398261, GQ398262, GQ398258, GQ298259) also changed between phylogenetic trees inferred across the DENV2 genome, and these were also removed as possible recombinants (Fig S1).

#### East African DENV2 Cosmopolitan viruses endemically circulate, but there is evidence of transmissions between East Africa and Asia

Kenyan genomes from Malindi cases were found in a monophyletic, well supported cluster separated from other African and international genomes in the sub-lineage with a long branch (Fig 1), indicating national-scale spatial structure of Kenyan populations and endemic circulation. They were most closely related to whole genome viruses from Pakistan and India (Fig 1), suggestive of dispersal between South/South-West Asia and East Africa. However, this result must be cautiously interpreted given the long branch ancestral to the Kenyan cluster, consistent with the very limited whole genome sampling of DENV-2 in other African countries, and elsewhere in Kenya. The *env*-gene phylogeny contained more African sequences from Burkina Faso, Ghana, Tanzania, Uganda and Somalia improving the resolution of DENV-2 Cosmopolitan in this region (Fig 2). This phylogeny indicates that DENV-2 Kenya African viruses were most closely related to Indian strains (Fig 2), although the branch length of the Kenyan clade was still long even in the *env* tree. The sole Uganda strain also clustered with Indian sequences, but its poor branch support also includes the possibility of previous transmissions between Kenya and Uganda (Fig 2). This extended *env* analysis also indicated that the strains from the 2011 Somalia outbreak were not ancestral to the sampled 2017 Kenya strains. Indeed, these were positioned into an entirely separate clade which included strains from Burkina Faso and Ghana and which comprised an “East-West” African cluster sampled over 28 years, indicating long term endemic African circulation of the DENV-2 Cosmopolitan genotype. Interestingly, this East-West cluster suggested spread between Africa and Saudi Arabia (Fig 2). Asian-African transmission was also inferred in the well supported clustering of a Tanzanian strain in a Chinese clade (Fig 2).

**Figure 2.**
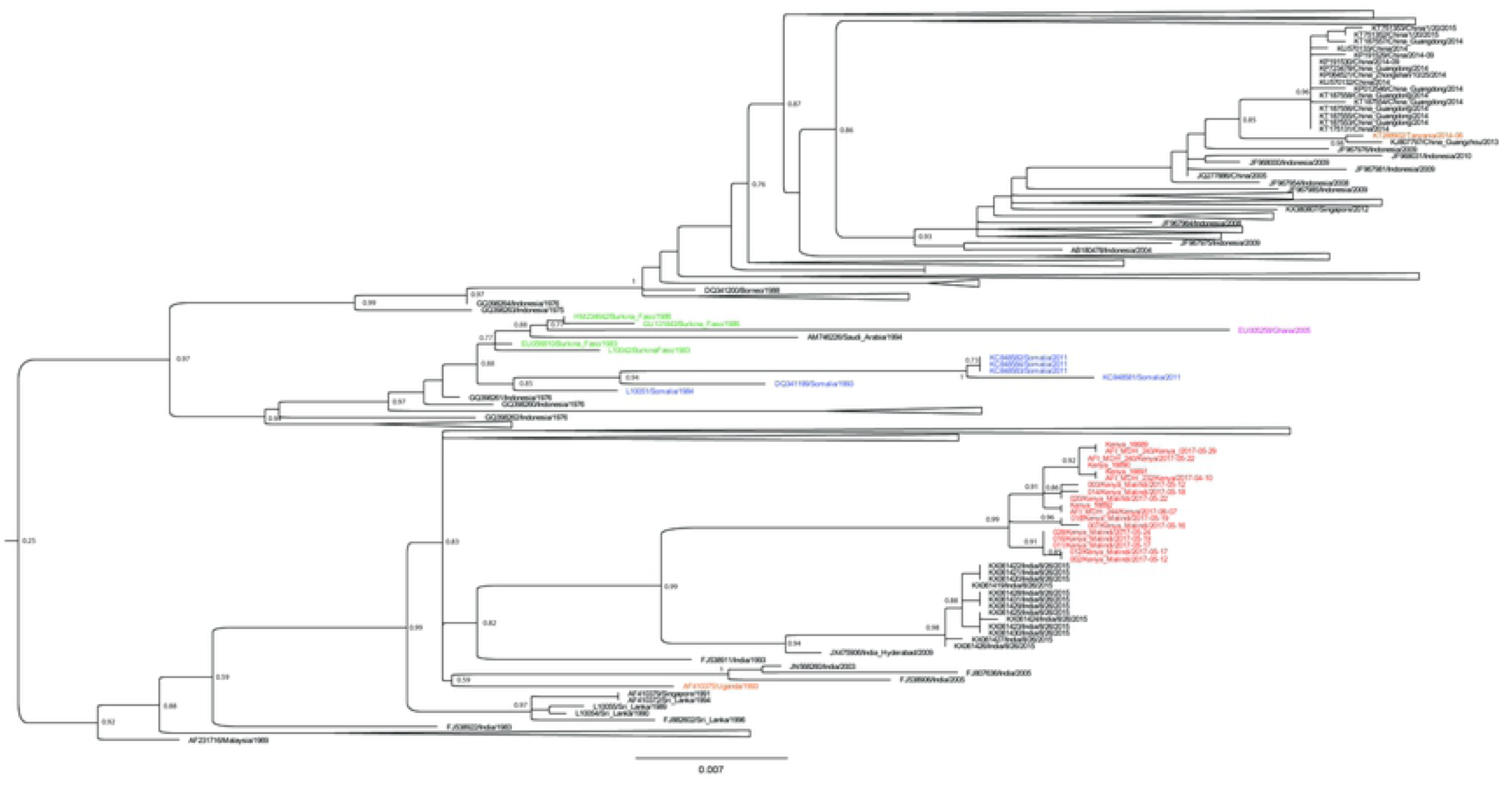
Maximum likelihood phylogeny of African *env* sequence data and reference DENV-2 Cosmopolitan *env* sequence data. Some background data clades collapsed to improve clarity, and all taxa names are made available in full with the removed American genotype outgroup in supplemental Figure S2. Data are color-coded by African country (Kenya = red, Uganda = orange, Burkina Faso = green, Ghana = purple, Somalia = blue, Tanzania = brown), and all non-African countries are represented by black. Scale bar indicates genetic distance (substitutions/site).

Given the regional importance of this “East-West” African cluster, we performed an extended time-scaled Bayesian analysis in BEAST to estimate the geographic history of this endemic DENV-2 lineage, the methodological details of which are indicated in the supplemental Box S1. A maximum clade credibility tree confirmed that Burkina Faso seeded neighboring Ghana outbreaks in 2005, and indicated a high probability of transmission of DENV-2 from Africa into neighboring Saudi Arabia (Fig 3). This same analysis estimated that this lineage of DENV-2 emerged in Somalia and Burkina Faso as early as 1978 (TMCRA 1981, 95% HPD 1978-1983) and 1978 (TMRCA 1980, 95% HPD 1977-1982), respectively. The dated root the East-West cluster indicated emergence of this DENV-2 strain in Africa as early as 1975 (TMRCA 1978, 95% HPD 1975-1981). The exact origins of this strain remains unclear, with only moderate probability support for Burkina Faso as the country of emergence (Fig 3), suggesting that other unsampled African or non-African countries may have played a key role of the early epidemic history of this East-West sublineage.

**Figure 3.**
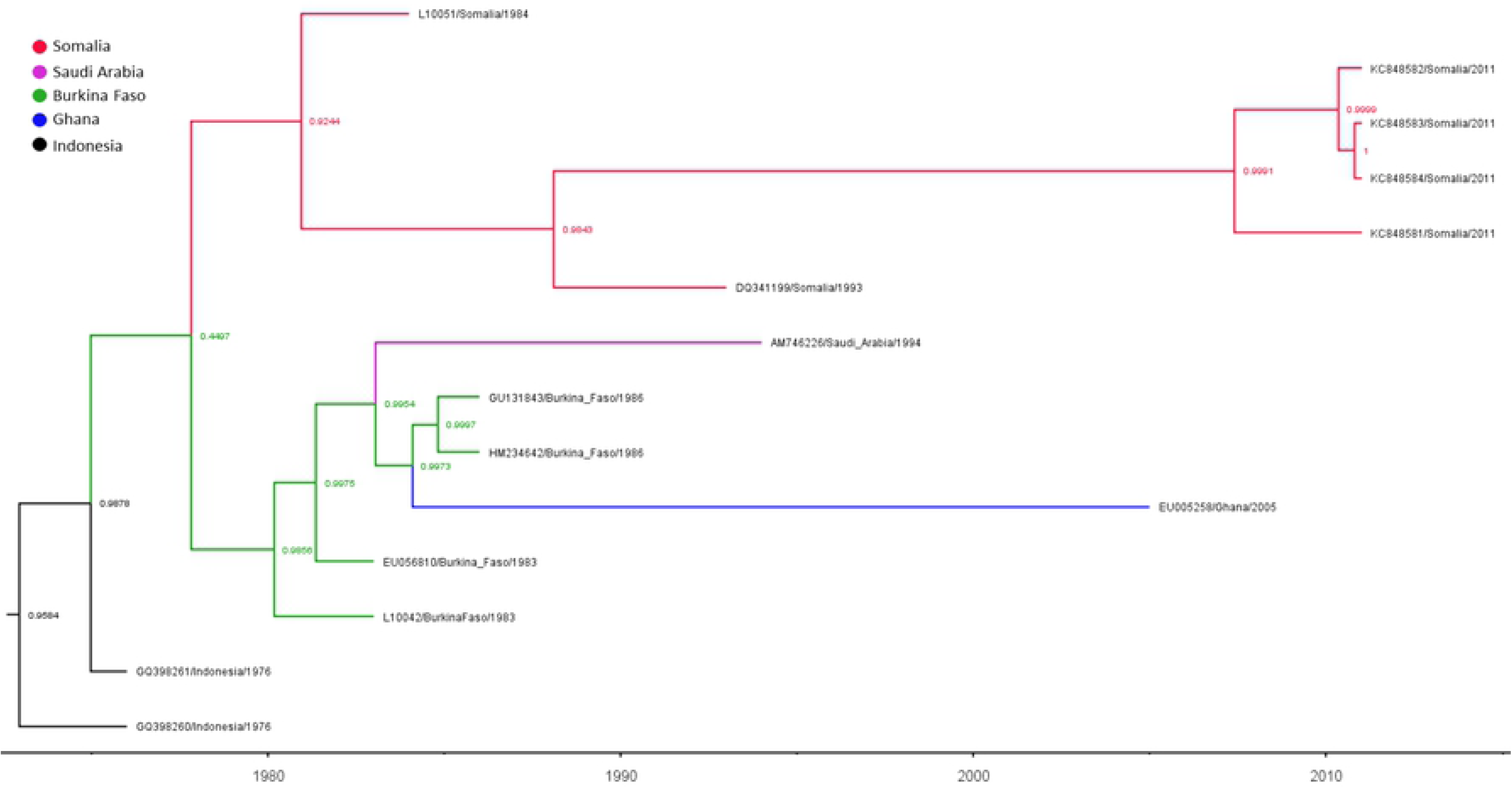
Bayesian time-scaled phylogeny of the Cosmopolitan ‘East-West’ African cluster. The scale represents time with scale of years. Color legend refers to the geographic location of collection of the sampled viruses (tips) as well as inferred ancestral strains. Numbers refer to the geographic state probability of nodes

#### The current 2016-2017 DENV2 outbreak into Malindi arose from strains introduced in 2013

There was weak evidence only (probability = 0.55) that the recent DENV-2 Cosmopolitan Kenyan clade was originally introduced from India, and this poor statistical support primarily reflects geographic and temporal sampling gaps in Africa (Fig 4). The time-to-most-common-recent-ancestor of the node which defined all Kenyan Malindi DENV-2 Cosmopolitan strains was estimated at September 2013 (95% credible interval January 2012 – May 2015), thereby indicating that this particular DENV clade has been circulating in Kenya for approximately 3.72 years (2.06 – 5.38 years) (Fig 4), although such an estimate is limited by a paucity of data sampled from other Kenyan locales. This Bayesian analysis estimated the mean evolutionary rate of the entire Cosmopolitan sub-lineage to be 8.94 × 10^−4^ subs/site/year (95% credible interval 5.7 × 10^−4^ to 1.28 x10^−3^), which is comparable to other estimates of DENV-2 evolutionary rates [7]. We further showed that there was considerable variability in strain evolutionary rate across this entire Cosmopolitan sub-lineage (coefficient of variation = 0.56, 95% credible interval 0.31 – 0.89) supporting a relaxed molecular clock model. Indeed, there was greater than three-fold variability in the evolutionary rates of the ancestral strains of the Kenyan outbreak alone (range 5 × 10^−4^ through to 1.8 × 10^−3^ substitutions/site/year). Analysis at this spatial scale also indicated that the Kenyan cluster separates into at least two well defined clades with strain co-circulation even at this fine spatial scale, although it is unclear if this represents two distinct introductions of two strains or *in-situ* diversification of a single introduced strain (Fig 4).

**Figure 4.**
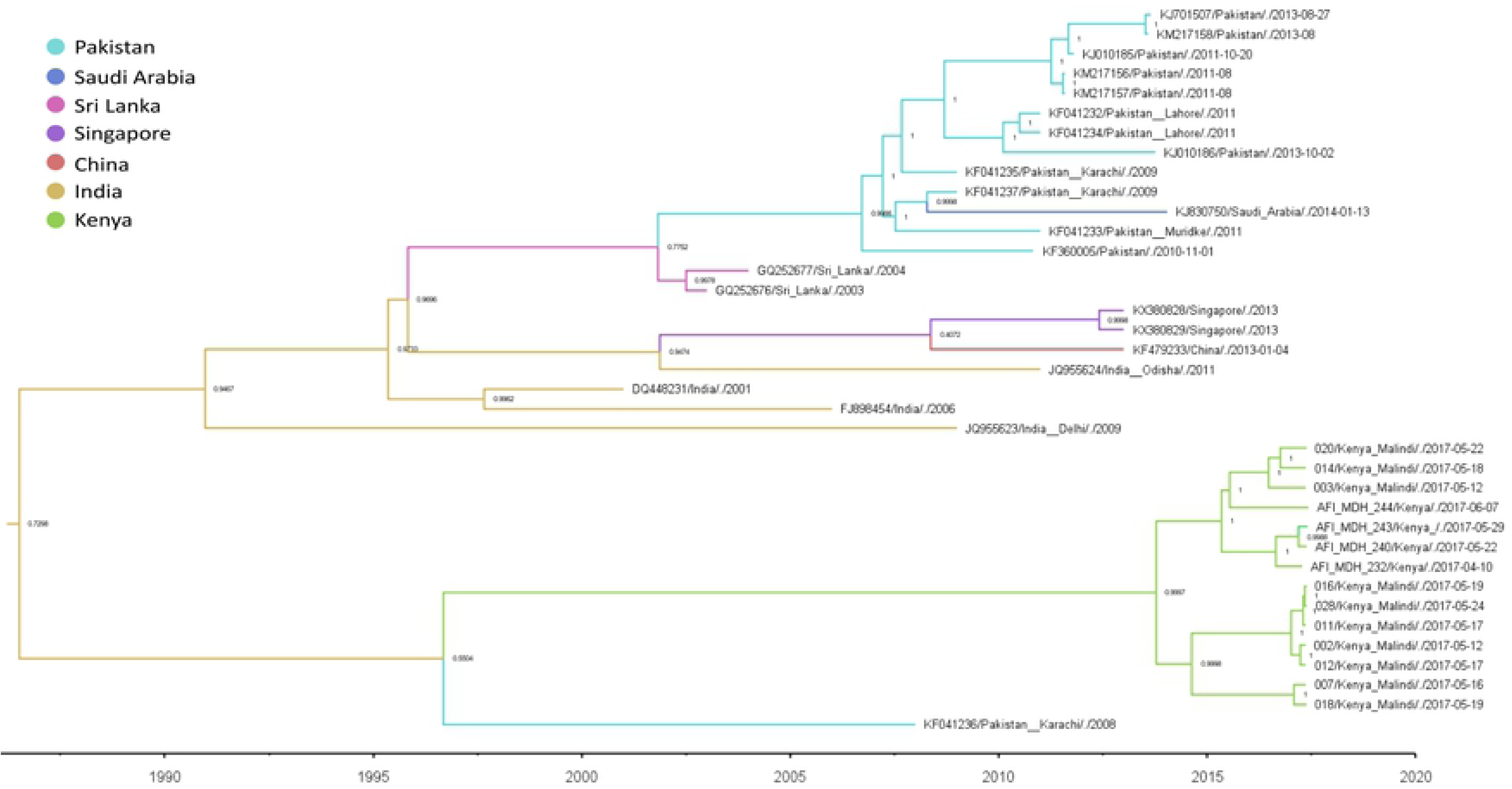
Bayesian time-scaled phylogeny of the Cosmopolitan sub-lineage containing all Kenya data. Time-scale not shown. Color legend refers to the geographic location of collection of sampled viruses (tips) as well as inferred ancestral strains. Numbers refer to the geographic state probability of nodes.

#### Positive selection and immune escape may explain epidemic diversification during the 2017 DENV-2 Kenyan outbreak

Table 1 shows the codons along the entire genome of the whole DENV-2 Cosmopolitan lineage which are estimated to be under positive selection, and most of these were found on the non-structural genes. Of these seven codon locations, there was strong evidence (positive by three methods) for diversifying selection pressure at codon 1867 which is located in the NS3 gene, and codon 2762 in the NS5 gene. In addition to the above analyses of the Cosmopolitan lineage as a whole, the FEL approach can be used to compare different selection pressures within specific sub-lineages or clades within the dataset (Table 2). Thus, Kenyan clade was selected to be used as the sub-dataset for selection analyses and compared to the rest of the Cosmopolitan lineage which was set as the background dataset. Whole genome selection pressure analyses of the Kenyan clade alone revealed presence of positive/diversifying selection that was acting on the Kenyan genomes only (Table 2), although this finding is derived from one selection pressure method only.

**Table 1.** DENV-2 Cosmopolitan lineage codons estimated to be under diversifying (positive) selection pressure*

| Genome<br>codon<br>position | Protein | Statistical significance of positive<br>selection pressure detected |  |  | Comments |
| --- | --- | --- | --- | --- | --- |
|  |  | FUBAR<br>(probability) | SLAC<br>(p-value) | FEL<br>(p-value) |  |
| 171 | prM | 0.9167 | p > 0.1 | p = 0.054 | Weak evidence for positive selection, codon variation is not unique to Kenyan data. |
| 670 | Env | 0.9186 | p > 0.1 | p = 0.041 | Moderate evidence for positive selection, codon variation is not unique to Kenyan data. |
| 1867 | NS3 | 0.99 | p = 0.03 | p = 0.006 | Strong evidence of positive selection, found within CD8+ epitope predicted epitope<br>TFDSEYIKT, codon variation not unique to Kenyan data. |
| 2387 | NS4B | 0.97 | p > 0.1 | p = 0.016 | Moderate evidence for positive selection, codon variation not unique Kenyan data but<br>appears to have driven the diversification of the 2012-2013 DENV-2 Cosmopolitan<br>Singapore outbreak. |
| 2391 | NS4B | 0.98 | p > 0.1 | p = 0.049 | Moderate evidence for positive selection, codon variation not unique to Kenyan data but<br>appears to have driven the diversification of the 2012-2013 DENV-2 Cosmopolitan<br>Singapore outbreak. |
| 2762 | NS5 | 0.94 | $p = 0.04$ | $p = 0.038$ | Strong evidence for positive selection, codon variation is not unique to Kenyan data. |
| 3135 | NS5 | 0.92 | $p > 0.1$ | $p = 0.058$ | Weak evidence for positive selection, codon variation is not unique to Kenyan data. |

\*Codons found to be under positive selection by at least two methods only presented. Dataset included an American genotype outgroup strain.

**Table 2.**
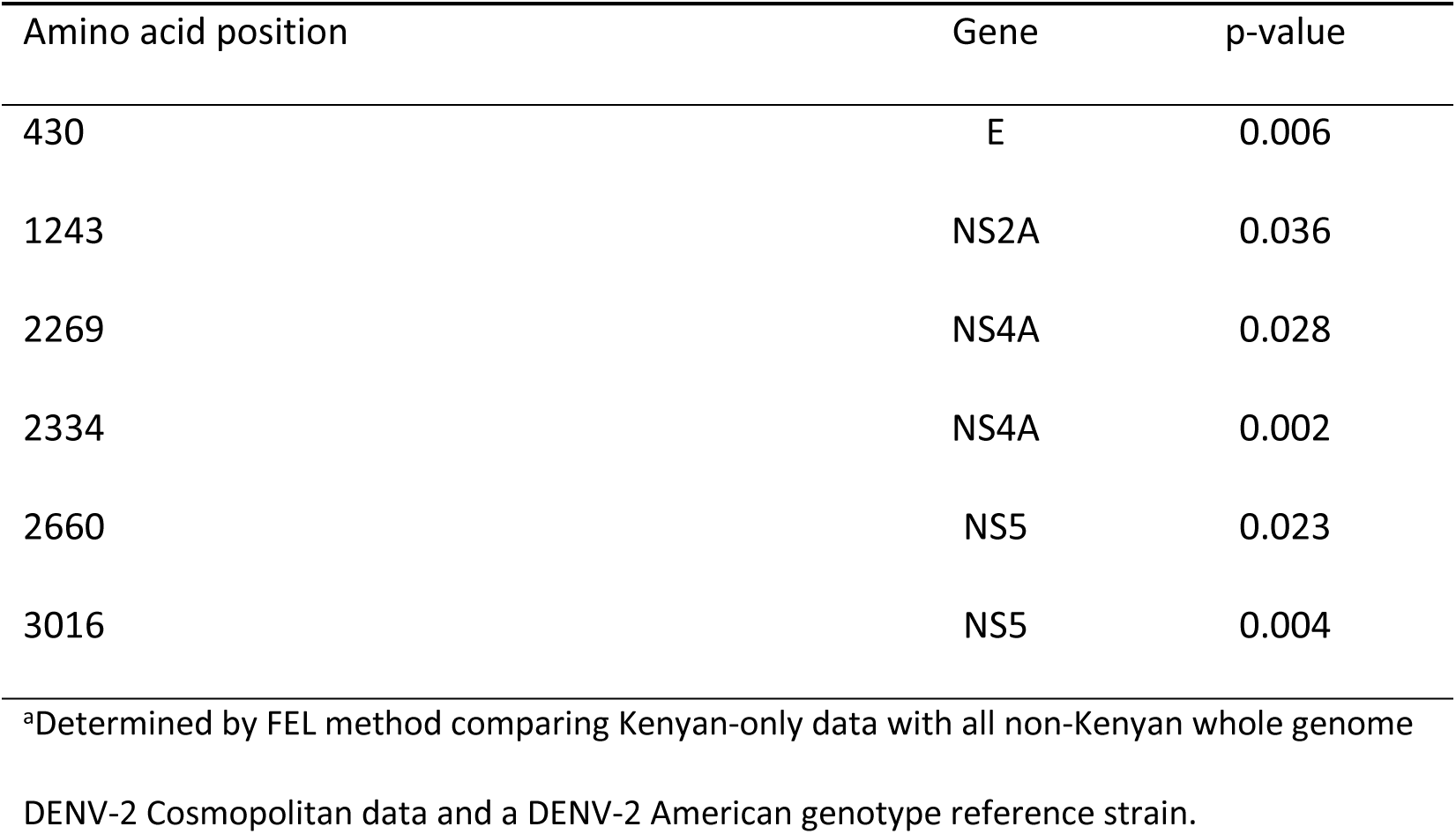
Amino acid positions under diversifying selection specific for the Kenyan strains sampled^a^

#### Extended Kenyan genome data reveals a possible novel MHC-I CD8+ epitope within the DENV-2 Cosmopolitan lineage

Through our CTL prediction analyses, we identified seven possible CTL epitopes across the Kenyan data (Table 3), the majority across the non-structural proteins. Only three of these have had previous experimental data to support their role as possible CD8+ CTL epitopes and/or CD4+ T-helper cell epitopes (FRKEIGRML, GWGNGCGLF and FTMRLLSPV). While the FRKEIGRML and GWGNGCGLF peptides are highly conserved across all existing published global DENV-2 Cosmopolitan whole genome data, FTMRLLSPV appears to be more geographically restricted to DENV viral populations in South Asia, South West Asia and Kenya (Table 3). Of the four potential novel epitopes, KMDIGAPLL was not found in any sampled non-Kenyan viral populations suggesting that there may be unique DENV CTL epitopes in those strains circulating Kenya. A position within the TFDSEYIKT epitope was found to be under positive selection for the Cosmopolitan lineage as a whole (Table 1), suggesting that this may be a substantial immunogen. Interestingly, one of the predicted CD8+ epitopes, FRKEIGRML in the capsid region, was found within a linear epitope associated with MHC-II positive assays, suggesting that this specific region may have high immunogenicity for development of both CD4+ and CTL responses to DENV2.

**Table 3.**
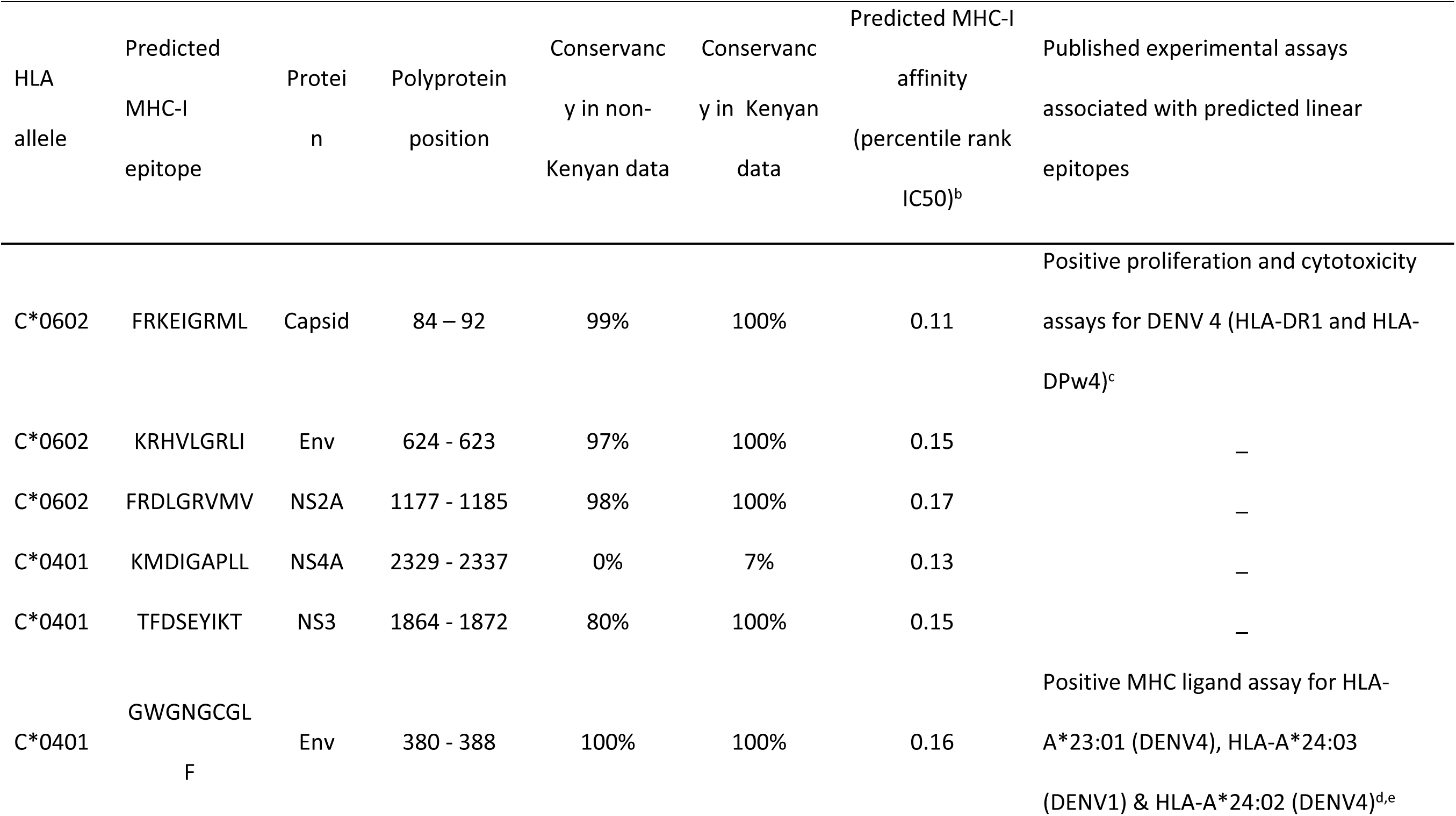
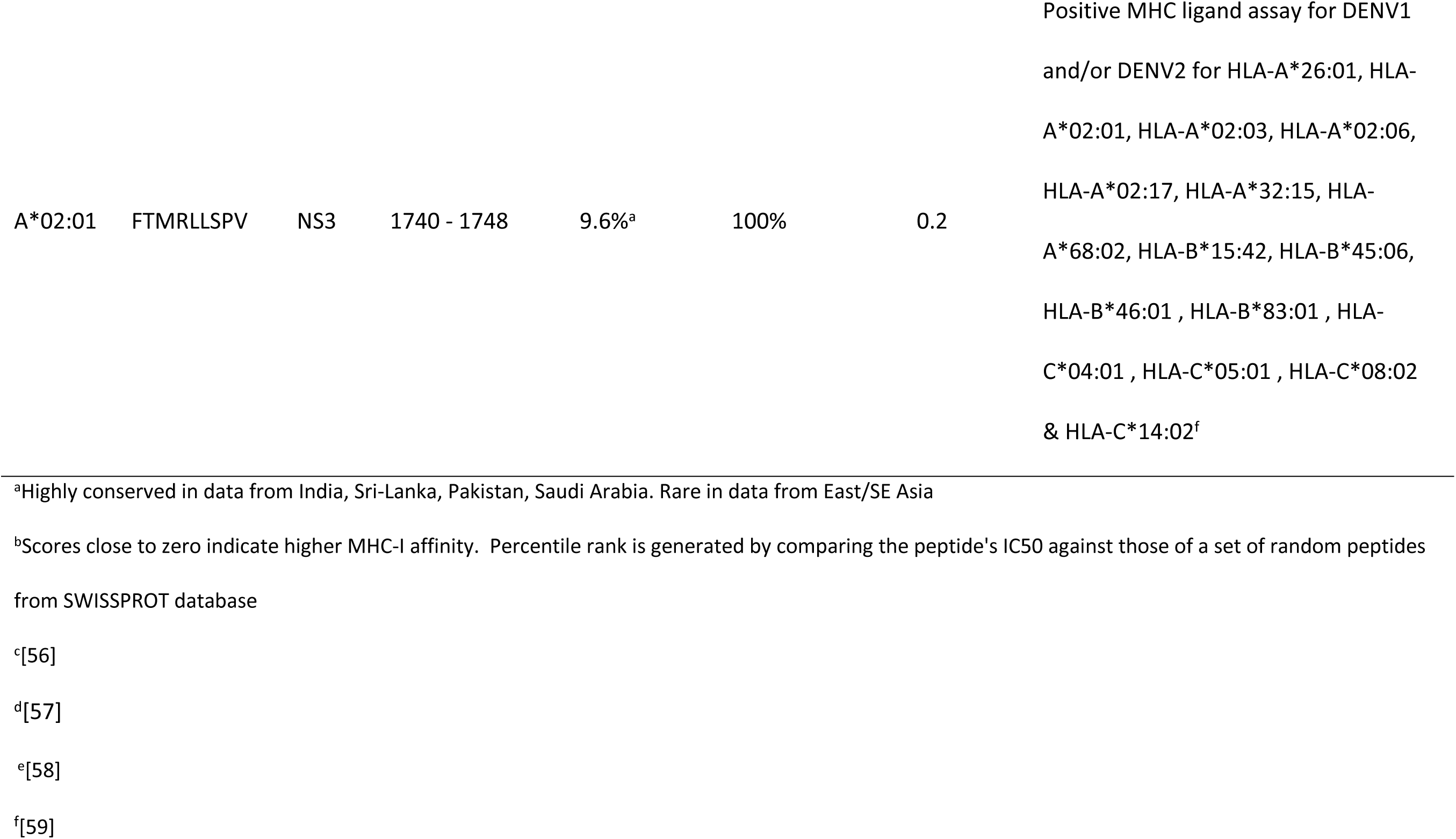
MHC-I epitopes predicted in Kenyan DENV-2 whole genome data

## Discussion

Prior to this study, there were a very limited number DENV-2 whole genomes were available from the entire continent of Africa. Our study enabled insights into the spatial structure of DENV-2 viruses in Africa. We show transmissions between Africa and Asia, as well as evidence of long term circulation of DENV-2 viruses in Africa with related lineages that span the geographic breadth of the continent and which may have emerged as early as the 1970s. Our analyses are in line with previous reports of Cosmopolitan lineage being the only DENV2 lineage found in Africa to date, however this may simply reflect under-sampling of DENV-2 in Africa. Indeed, the apparent sole circulation of the Cosmopolitan genotype is unusual given that other DENV-2 endemic continents, such as Asia, have experienced up to four other genotypes [31]. A study recently reported a DENV-2 Asian-II genotype strain isolated from West Africa, however the implausible lack of evolution for its sampling date rather indicates that this sample actually represents a New Guinea C 1944 contaminant [52].

We showed spatial structure to African DENV-2 epidemics on several scales. On an intercontinental scale we show evidence of DENV-2 dispersal between East Africa and proximal South-West Asia (Saudi Arabia). Within Africa, there was evidence of DENV-2 spread on an East-West axis, and high probability of spread between neighboring Burkina Faso and Ghana. Interestingly, our analysis suggests that the recent Kenyan 2017 DENV-2 outbreak was unrelated to the 2011 DENV outbreak in neighboring Somalia, highlighting the utility of genomic epidemiology in evaluating the origins of DENV outbreaks. Time-scaled genealogies indicate that DENV-2 viruses are also spatially structured within Kenya, with a time-to-most-common-recent ancestor analysis indicating that DENV-2 strains were circulating for up to 5.38 years in Kenya before detection in the 2017 Malindi outbreak, although such estimates are subject to unsampled viruses from other Kenyan locations. This estimate of pre-outbreak circulation time highlights that DENV strains may circulate in a population for years before detection by sentinel surveillance mechanisms. Indeed, this has been shown in other DENV endemic tropical countries, such as Singapore [53]. Our analyses also indicated that there was spatial structure of dengue viruses at a subnational scale, with two co-circulating clades detected within Malindi. Such fine-scale co-circulation of within-genotype clades has also been noted in other tropical dengue-endemic regions such as Thailand [54]. A caveat to these spatial epidemiological conclusions is that they are susceptible to unsampled dengue viruses elsewhere in Kenya and neighboring regions. The influence of missing data on phylogeographic conclusions is well known [7], and further sampling in complementary genomic epidemiology studies from this country will be critical to clarify the epidemiology of DENV-2 in the Coastal Kenyan and greater East African region.

Selection pressure analyses indicated Kenyan DENV strains uniquely being under positive selection, predominantly across the nonstructural genes, which may reflect population immunity escape associated with the 2016-2017 outbreak viral variants, and which may predict epidemic diversification of these strains among other Kenyan populations. A caveat to this is that selection pressure analyses are best confirmed with a least three independent methods, yet such methods in this case do not allow the direct comparison of the Kenyan clade to background DENV-2 Cosmopolitan data and only offer estimation of selection pressure across the entire Cosmopolitan lineage (Table 1). Another caveat is that dengue virus selection pressure analyses are also inherently susceptible to the size and sampling distribution of the genomic datasets analyzed [7], and our findings here should be compared and contrasted with other studies sampling dengue viruses in Kenyan and other East African settings, ideally with similar methods.

Among the Kenyan sequence data, linear 9mer peptides which flanked the positively selected amino acids specific for the Kenyan genomes (Table 2) were not found in any known DENV linear CTL epitopes documented in IEDB [51]. If the detected positive selection in these positions was indeed due to population immunologic pressures, this would indicate that there might be yet undiscovered DENV epitopes and immunogenic regions specific for this part of the world. Interestingly, epitope prediction analyses showed that one of these lineage-wide positive selection sites resides within a predicted MHC-I CD8+ DENV-2 Cosmopolitan virus epitope only evident in a sampled Kenyan virus (Table 3), offering further evidence that there are viral-host interactions unique to DENV-2 viruses circulating in the Kenyan population. This should be confirmed with further DENV genome sampling, more precise T-cell epitope prediction using host HLA typing, and in-vitro cell-mediated immunity assays, particularly as such peptides may be important immunogens of relevance to dengue vaccine design and evaluation [55].

Taken together, our findings indicate that the 2017 Kenyan Malindi DENV-2 outbreak arose from a strain which had circulated for several years in Kenya before recent detection, has experienced diversifying selection pressure, and may contain new putative immunogens relevant to vaccine design. While limited by a relatively small number of absolute DENV-2 genomes, this study is an important step toward redressing our limited understanding of the virology and epidemiology of DENV viruses in this country, and more broadly in Africa, and should prompt further whole genome sequencing of dengue viruses in this and other African countries.

## Acknowledgements

The authors are grateful for those research subjects who participated in this study.

## Disclaimer

The views expressed in this article are those of the authors and do not necessarily reflect the official policy or position of the Department of the Army, Department of Defense nor the U.S. Government. Several of the authors are US Government Employees. This work was prepared as part of their official duties. Title 17 U.S.C. § 105 provides that ‘Copyright protection under this title is not available for any work of the United States Government.’ Title 17 U.S.C. §101 defines a U.S. Government work as a work prepared by a military service member or employee of the U.S. Government as part of that person’s official duties

## Supporting Information Legends

**Figure S1.** Neighbor-joining trees on four sub-genomic regions (CDS nucleotide positions 1-2543, 2544-5087, 5088-7630 and 7631-10173). Scale refers to nucleotide substitutions/site.

**Figure S2.** Maximum likelihood phylogeny of full DENV-2 Cosmpolitan env genes. American genotype outgroup removed for clarity. Scale refers to nucleotide substitutions/site. Numbers refer to aLRT values. Taxa tables are right-aligned for clarity, and are color-coded by African country (Kenya = red, Uganda = orange, Burkina Faso = green, Ghana = purple, Somalia = blue, Tanzania = brown). All non-African taxa labels are indicated in black.

## Supplemental material (extended)

Additional methods, primers, and reference GenBank accession numbers.

